# The Genetic Legacy of the Expansion of Turkic-Speaking Nomads Across Eurasia

**DOI:** 10.1101/005850

**Authors:** Bayazit Yunusbayev, Mait Metspalu, Ene Metspalu, Albert Valeev, Sergei Litvinov, Ruslan Valiev, Vita Akhmetova, Elena Balanovska, Oleg Balanovsky, Shahlo Turdikulova, Dilbar Dalimova, Pagbajabyn Nymadawa, Ardeshir Bahmanimehr, Hovhannes Sahakyan, Kristiina Tambets, Sardana Fedorova, Nikolay Barashkov, Irina Khidiatova, Evelin Mihailov, Rita Khusainova, Larisa Damba, Miroslava Derenko, Boris Malyarchuk, Ludmila Osipova, Mikhail Voevoda, Levon Yepiskoposyan, Toomas Kivisild, Elza Khusnutdinova, Richard Villems

**Author notes:** These authors contributed equally (Bayazit Yunusbayev, Mait Metspalu). Corresponding author: Bayazit Yunusbayev; Evolutionary Biology group, Estonian Biocentre, Tartu 51010, Estonia.

## Abstract

The Turkic peoples represent a diverse collection of ethnic groups defined by the Turkic languages. These groups have dispersed across a vast area, including Siberia, Northwest China, Central Asia, East Europe, the Caucasus, Anatolia, the Middle East, and Afghanistan. The origin and early dispersal history of the Turkic peoples is disputed, with candidates for their ancient homeland ranging from the Transcaspian steppe to Manchuria in Northeast Asia. Previous genetic studies have not identified a clear-cut unifying genetic signal for the Turkic peoples, which lends support for language replacement rather than demic diffusion as the model for the Turkic language’s expansion. We addressed the genetic origin of 373 individuals from 22 Turkic-speaking populations, representing their current geographic range, by analyzing genome-wide high-density genotype data. Most of the Turkic peoples studied, except those in Central Asia, genetically resembled their geographic neighbors, in agreement with the elite dominance model of language expansion. However, western Turkic peoples sampled across West Eurasia shared an excess of long chromosomal tracts that are identical by descent (IBD) with populations from present-day South Siberia and Mongolia (SSM), an area where historians center a series of early Turkic and non-Turkic steppe polities. The observed excess of long chromosomal tracts IBD (> 1cM) between populations from SSM and Turkic peoples across West Eurasia was statistically significant. Finally, we used the ALDER method and inferred admixture dates (∼9th–17th centuries) that overlap with the Turkic migrations of the 5th–16th centuries. Thus, our results indicate historical admixture among Turkic peoples, and the recent shared ancestry with modern populations in SSM supports one of the hypothesized homelands for their nomadic Turkic and related Mongolic ancestors.

**Author Summary:** Centuries of nomadic migrations have ultimately resulted in the distribution of Turkic languages over a large area ranging from Siberia, across Central Asia to Eastern Europe and the Middle East. Despite the profound cultural impact left by these nomadic peoples, little is known about their prehistoric origins. Moreover, because contemporary Turkic speakers tend to genetically resemble their geographic neighbors, it is not clear whether their nomadic ancestors left an identifiable genetic trace. In this study, we show that Turkic-speaking peoples sampled across the Middle East, Caucasus, East Europe, and Central Asia share varying proportions of Asian ancestry that originate in a single area, southern Siberia and Mongolia. Mongolic- and Turkic-speaking populations from this area bear an unusually high number of long chromosomal tracts that are identical by descent with Turkic peoples from across west Eurasia. Admixture induced linkage disequilibrium decay across chromosomes in these populations indicates that admixture occurred during the 9th–17th centuries, in agreement with the historically recorded Turkic nomadic migrations and later Mongol expansion. Thus, our findings reveal genetic traces of recent large-scale nomadic migrations and map their source to a previously hypothesized area of Mongolia and southern Siberia.

## Introduction

Linguistic relatedness is frequently used to inform genetic studies [1] and here we take this path to reconstruct aspects of a major and relatively recent demographic event, the expansion of nomadic Turkic-speaking peoples, who reshaped much of the West Eurasian ethno-linguistic landscape in the last two millennia. Modern Turkic-speaking populations are a largely settled people; they number over 170 million across Eurasia and, following a period of migrations spanning the ∼5th–16th centuries, have a wide geographic dispersal, encompassing Eastern Europe, Middle East, Northern Caucasus, Central Asia, Southern Siberia, Northern China, and Northeastern Siberia [2-4].

The extant variety of Turkic languages spoken over this vast geographic span reflects only the recent (2100–2300 years) history of divergence, which includes a major split into Oghur (or Bolgar) and Common Turkic [5,6]. This period was preceded by early Ancient Turkic, for which there is no historical data, and a long-lasting proto-Turkic stage, provided there was a Turkic-Mongolian linguistic unity (protolanguage) around 4500–4000 BCE [7,8].

The earliest Turkic ruled polities (between the 6th and 9th centuries) were centered in what is now Mongolia, northern China, and southern Siberia. Accordingly, this region has been put forward as the point of origin for the dispersal of Turkic-speaking pastoral nomads [3,4]. We designate it here as an “Inner Asian Homeland” (IAH) and note at least two issues with this working hypothesis. First, the same approximate area was earlier dominated by the Xiongnu Empire (Hsiung-nu) (200 BCE–100 CE) and later by the short-lived Xianbei (Hsien-pi) Confederation (100–200 CE) and Rouran State (aka Juan-juan or Asian Avar) (400–500 CE). These steppe polities were likely established by non-Turkic-speaking peoples and presumably united ethnically diverse tribes. It is only in the second half of the 6th century that Turkic-speaking peoples gained control of the region and formed the rapidly expanding Göktürk Khaganate, succeeded soon by numerous khanates and khaganates extending from northeastern China to the Pontocaspian steppes in Europe [2-4]. Secondly, Göktürks represent the earliest known ethnic unit whereby Turkic peoples appear under the name Turk. Yet, Turkic-speaking peoples appear in written historical sources before that time, namely when Oghuric Turkic-speaking tribes appear in the Northern Pontic steppes in the 5th century, much earlier than the rise of Göktürk Khaganate in the IAH. Thus, the early stages of Turkic dispersal remain poorly understood and our knowledge about their ancient habitat remains a working hypothesis.

Previous studies based on Y chromosome, mitochondrial DNA (mtDNA), and autosomal markers show that while the Turkic peoples from West Asia (Anatolian Turks and Azeris) and Eastern Europe (Gagauzes, Tatars, Chuvashes, and Bashkirs) are generally genetically similar to their geographic neighbors, they do display a minor share of both mtDNA and Y haplogroups otherwise characteristic of East Asia [9-14]. Expectedly, the Central Asian Turkic speakers (Kyrgyz, Kazakhs, Uzbeks, and Turkmens), share more of their uniparental gene pool (9–76% of Y chromosome and over 30% of mtDNA lineages) with East Asian and Siberian populations [15,16]. In this regard, they differ from their southern non-Turkic neighbors, including Tajiks, Iranians, and different ethnic groups in Pakistan, except Hazara. However, these studies do not aim to identify the precise geographic source and the time of arrival or admixture of the East Eurasian genes among the contemporary Turkic-speaking peoples. The “eastern” mtDNA and even more so Y-chromosome lineages (given the resolution available to the studies at the time) lack the geographic specificity to explicitly distinguish between regions within Northeast Asia and Siberia, and/or Turkic and non-Turkic speakers of the region [17,18].

Several studies using genome-wide SNP panel data describe the genetic structure of populations in Eurasia and although some include different Turkic populations [14,19-22], they do not focus on elucidating the demographic past of the Turkic-speaking continuum. In cases where more than one geographic neighbor is available for comparison, Turkic-speaking peoples are genetically close to their non-Turkic geographic neighbors in Anatolia [21,23], the Caucasus [14], and Siberia [20,22]. A recent survey of worldwide populations revealed a recent (13th–14th century) admixture signal among the three Turkic populations (Turks, Uzbeks, and Uygurs) and one non-Turkic population (Lezgins) with Mongolas (from northern China), the Daurs (speaking Mongolic language), and Hazaras (of Mongol origin) [24]. This study also showed evidence for admixture (dating to the pre-Mongol period of 440–1080 CE) among non-Turkic (except Chuvashes) East European and Balkan populations with the source group related to modern Oroqens, Mongolas, and Yakuts. This is the first genetic evidence of historical gene flow from a North Chinese and Siberian source into some north and central Eurasian populations, but it is not clear whether this admixture signal applies to other Turkic populations across West Eurasia.

Here we ask whether it is possible to identify explicit genetic signal(s) shared by all Turkic peoples that have likely descended from putative prehistoric nomadic Turks. Specifically, we test whether different Turkic peoples share genetic heritage that can be traced back to the hypothesized IAH. More specifically, we ask whether this shared ancestry occurred within an historical time frame, testified by an excess of long chromosomal tracts identical by descent between Turkic-speaking peoples across West Eurasia and those inhabiting the IAH. To address these questions we used a genome-wide high-density genotyping array to generate data on Turkic-speaking peoples representing all major branches of the language family (Figure 1B).

**Figure 1.**
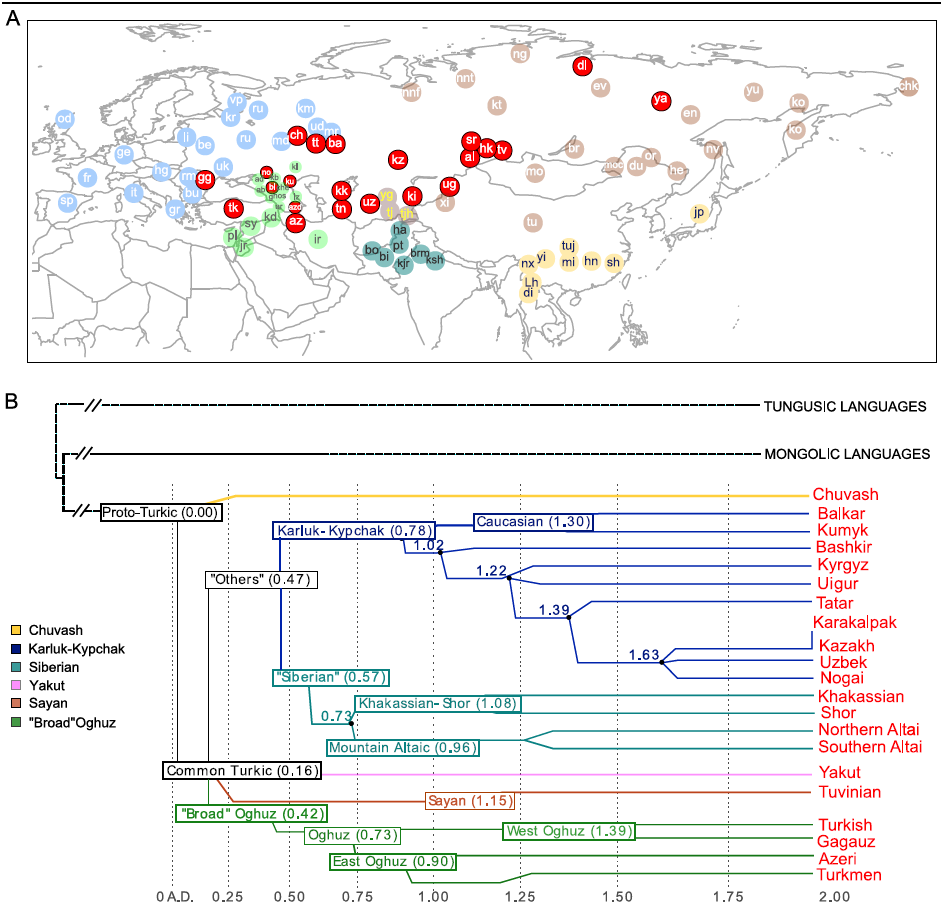
**Geographic map of samples included in this study and linguistic tree of Turkic languages.** Panel A) Non-Turkic-speaking populations are shown with light blue, light green, dark green, light brown, and yellow circles, depending on the region. Turkic-speaking populations are shown with red circles regardless of the region of sampling. Full population names are given in Table S1, Panel B) The linguistic tree of Turkic languages is adapted from Dybo 2004 and includes only those languages spoken by the Turkic peoples analyzed in this study. The x-axis shows the time scale in kilo-years (kya). Internal branches are shown with different colors.

### Results

To characterize the population structure of Turkic-speaking populations in the context of their geographic neighbors across Eurasia, we genotyped 322 new samples from 38 Eurasian populations and combined it with previously published data (see Table S1 and Material and Methods for details) to yield a total dataset of 1,444 samples genotyped at 515,841 markers. The novel samples introduced in this study geographically cover previously underrepresented regions like Eastern Europe (Volga-Ural region), Central Asia, Siberia, and the Middle East. We used a *STRUCTURE*-like [25] approach implemented in the program ADMIXTURE [26] to explore the genetic structure in the Eurasian populations by inferring the most likely number of genetic clusters and mixing proportions consistent with the observed genotype data (from *K* = 3 through *K* = 14 groups) (Figure S1). As shown in previous studies [14,19,27] East Asian populations commonly contained alleles that find membership in two general clusters, shown here as k6 and k8, in a model assuming *K* = 8 “ancestral” populations (Figure 2). Geographically, the spread zones of these two components (clusters) were centered on Siberia and East Asia, respectively. Their combined prevalence declined as one moves west from East Asia. Overall, alleles from the Turkic populations sampled across West Eurasia showed membership in the same set of West Eurasian genetic clusters, k1–k4, as did their geographic neighbors. In addition, the Volga-Uralic Turkic peoples (Chuvashes, Tatars, and Bashkirs) also displayed membership in the k5 cluster, which contained the Siberian Uralic-speaking populations (Nganasans and Nenets) and extended to some of the European Uralic speakers (Maris, Udmurts, and Komis). However, in most cases the Turkic peoples showed a higher combined presence of the “eastern components” k6 and k8 than did their geographic neighbors.

**Figure 2.**
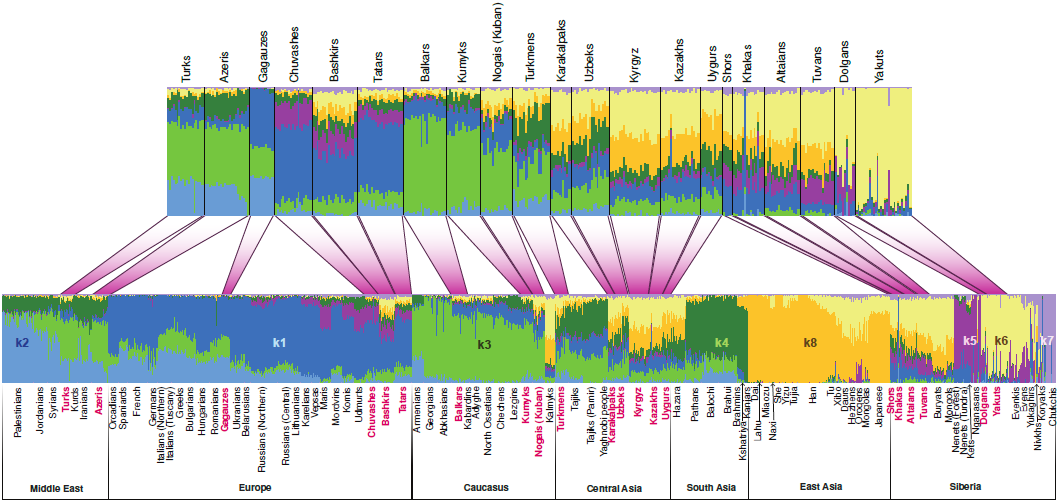
**Population structure inferred using ADMIXTURE analysis.** ADMIXTURE results at *K* = 8 are shown. Each individual is represented by a vertical (100%) stacked column indicating the proportions of ancestry in K constructed ancestral populations. Turkic-speaking populations are shown in red. The upper barplot shows only Turkic-speaking populations.

### Geographic distribution of recent shared ancestry

A recent study shows that even a pair of unrelated individuals from the opposite ends of Europe share hundreds of chromosomal tracts of IBD from common ancestors that lived over the past 3,000 years. The amount of such recent ancestry declines exponentially with geographic distance between population pairs, and such an isolation-by-distance pattern can be distorted due to population expansion or gene flow [28]. We observed a reasonably high correlation (correlation coefficient = 0.83, *p* < 0.0001) between the rate of IBD sharing decay and geographic distance when we removed the West Eurasian Turkic populations (sampled in the Middle East, Caucasus, Eastern Europe, and Central Asia) from our dataset. This implies that IBD sharing in our set of Eurasian populations is largely explained by isolation by distance. However, when we included the western Turkic populations, the correlation between IBD sharing decay and geographic distance was weaker (0.77, *p* < 0.0001). This implies that IBD sharing between the Turkic samples and other populations departs slightly from the isolation-by-distance pattern. To identify populations for which IBD sharing with Turkic populations departs from an isolation-by-distance pattern, we first computed IBD sharing (the average length of genome IBD measured in centiMorgans) for each of the 12 western Turkic populations with all other populations in the dataset (Table S2) and then subtracted the same statistic computed for their geographic neighbors (see the Materials and Methods section for details and Figure S2 for a schematic representation of this analysis). When the differences were overlaid for all 12 Turkic populations, we detected an unusually high signal of accumulated IBD sharing (samples indicated by a “plus symbol” on Figure 3 A, B, and C) for populations outside West Eurasia. The correlated signal of IBD sharing for these distant populations exceeded the expectation based on an isolation-by-distance model. Most of these distant populations are located in South Siberia and Mongolia (SSM) and Northeast Siberia, except the two samples in Eastern Europe (Maris) and the North Caucasus (Kalmyks). We note that the null hypothesis for this analysis assumed no systematic difference between any of the Turkic populations and their respective geographic neighbors. Therefore, the null hypothesis predicted that random differences accumulated across the entire geographic range of the western Turkic populations. To demonstrate this null expectation, we replaced each of the western Turkic populations by populations randomly drawn from the sets of respective non-Turkic neighbors, and repeated this subtraction/accumulation analysis, as shown in Figure S2. When the sets of random non-Turkic samples were tested, the accumulated signal was restricted to populations (indicated by the “plus symbol” on Figure S3) within West Eurasia, as expected by the null hypothesis. There are, however, two exceptions (Nganasans and Nenets) that, when examined closely, suggest an interesting finding consistent with our ADMIXTURE results. These two Siberian populations, Nganasans and Nenets (Figure S3 A, B, E, I, and J), speak Uralic languages and demonstrated a high accumulated signal only when our tested sets contained the western Uralic speakers (Maris, Komis, Vepsas, and Udmurts). This was in line with our ADMIXTURE results (Figure 2), as the k5 ancestry component was shared specifically between these western Uralic speakers and the two Siberian Uralic-speaking Nganasans and Nenets. We now return to the overall difference between the accumulated IBD sharing signal under the null hypothesis (see Figure S3) and that observed for the set of western Turkic populations (Figure 3). Some of the populations in SSM and Northeast Siberia demonstrated a strong IBD sharing signal with the western Turkic populations and this pattern most likely indicates recent gene flow from Siberia. To narrow down the source of this gene flow it is important to know which of the Siberian populations are indigenous to their current locations. We show in the Discussion section that only Tuvans, Buryats, and Mongols from the SSM area are indigenous to their current locations (at least within the known historical time) and therefore this area is the best candidate for the source of recent gene flow into the western Turkic populations.

**Figure 3.**
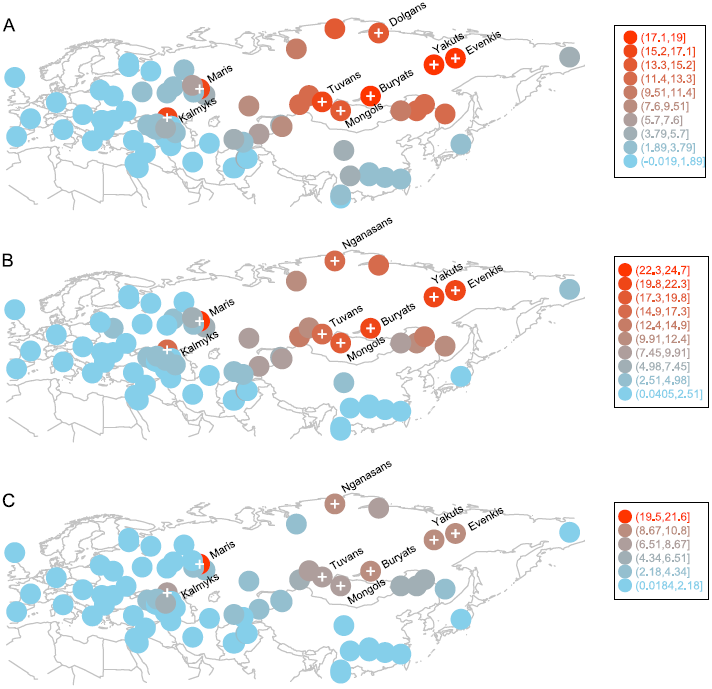
**Populations with high and correlated signals of IBD sharing with western Turkic peoples.** Circle positions correspond to population locations. Circle color indicates the amount of excess IBD sharing (shown in Legend) that a population shares with all 12 western Turkic populations. Populations with IBD sharing exceeding the 0.90 quantile are shown with a “plus symbol”. Panel A) IBD sharing signal based on IBD tracts of 1–2 cM. Panel B) IBD sharing signal based on IBD tracts of 2–3 cM. Panel C) IBD sharing signal based on IBD tracts of 3–4 cM

Our previous analysis suggests that the western Turkic populations (Table S2) experienced stronger gene flow from the SSM area than their non-Turkic neighbors, but it is not clear whether this signal is statistically significant. To test this, we computed IBD sharing between the group of SSM populations (Tuvans, Mongols, and Buryats, as well as a known migrant population, Evenkis) and each of the western Turkic populations. Then, for each of the western Turkic populations, we pooled their non-Turkic neighbors, and generated 10,000 permuted samples to see whether a comparable amount of IBD sharing (observed in tested Turkic populations) with the four Siberian populations is obtained by chance. IBD sharing was estimated separately for different classes of chromosomal tracts (1–2 cM, 2–3 cM, 3–4 cM, etc.), and permutation tests were performed. In most of the cases, higher IBD sharing between the western Turkic populations (compared to non-Turkic neighbors) and the Siberian populations was statistically significant (Figure 4 and Figure S4; numbers in red show how many Siberian populations have *p*-values ≤ 0.01). Some of the non-Turkic neighbors, such as Romanians, Lezgins, and North Ossetians, also shared a relatively high number of IBD tracts (Figure 4 and Figure S4) with the SSM populations. We conclude that the recent gene flow from the SSM area inferred in our previous analysis was not restricted to the western Turkic peoples, and the higher IBD sharing is evidence that Turkic populations are distinct from their non-Turkic neighbors.

**Figure 4.**
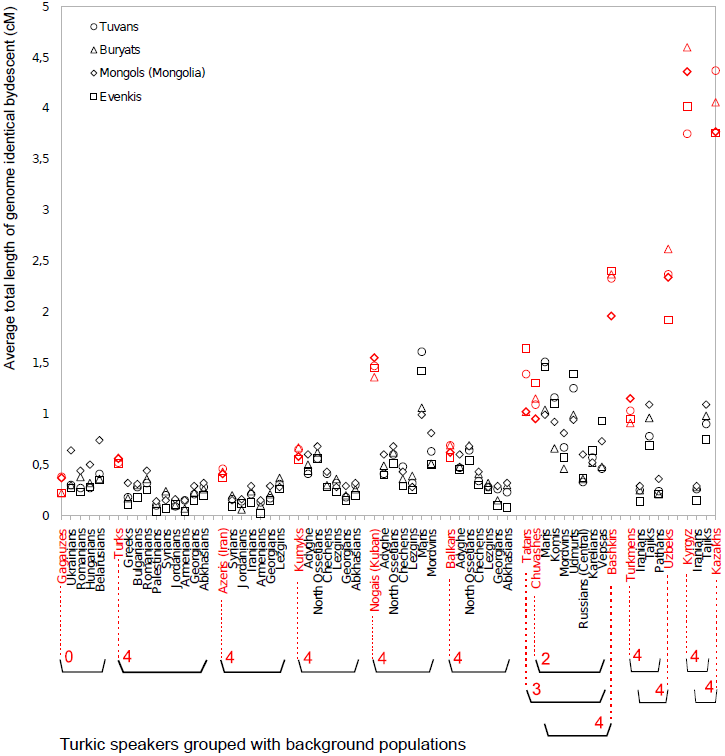
**Pairwise IBD sharing based on 1–2 cM long segments.** For each population ordered along the x–axis, IBD sharing is computed with three SSM populations (Tuvans, Buryats, Mongols) and Evenkis. Each Turkic-speaking population (shown in red) is grouped with its respective geographic neighbors using parentheses. The grouped geographic neighbors were pooled and used to perform a permutation test as described in the M&M section. Red numbers under the Turkic population name indicate how many SSM populations demonstrate a statistically significant excess of IBD sharing with a given Turkic population. Note that, for example, Bashkirs, Tatars, and Chuvashes share their geographic neighbors.

A spatial pattern in IBD sharing was noted when IBD tracts of different length classes were considered separately. For segment classes of 1–2 cM and 2–3 cM, higher IBD sharing is statistically significant for most Turkic speakers, except Gagauzes and Chuvashes (and Tatars in the case of 2–3 cM). For longer IBD tracts of 3–4 cM, statistical evidence for higher IBD sharing becomes weaker in some Middle Eastern and Caucasus (Azeris, Kumyks, and Balkars) samples. By weaker evidence, we mean that a statistically significant excess of IBD sharing was restricted to a subset of the four candidate ancestors tested. In the Volga-Ural region, for the same class of segments (3–4 cM), only Bashkirs continued to show strong evidence for gene flow, while Tatars and Chuvashes do not. For these two Turkic populations, not all tests were statistically significant because the background group, from which permuted samples are drawn, contained a Finnic speaking Mari people showing comparable levels of Asian admixture (Figure 2) and IBD sharing (Figure S4). When we considered even longer segments (4–5 cM and 5–6 cM), we no longer observed a systematic excess of IBD sharing for Turkic peoples in the Middle East, the Caucasus, or in the Volga-Ural region. In contrast, populations closer to the SSM area (Uzbeks, Kazakhs, Kyrgyz, and Uygurs, and also Bashkirs from the Volga-Ural region) still demonstrated a statistically significant excess of IBD sharing. This spatial pattern can be partly explained by a relative rarity of longer IBD tracts compared to shorter ones and recurrent gene flow events into populations closer to the SSM area.

### Dating the age of Asian admixture using the ALDER and SPCO methods

According to historical records, the Turkic migrations took place largely during ∼5th–16th centuries (little is known about earlier periods) and partly overlap with the Mongol expansion. Assuming 30 years per generation, the common Siberian ancestors of various Turkic peoples lived prior to and during this migration period between 20 and 53 generations ago. The expected length of a single-path IBD tract inherited from a common ancestor that lived ∼20–53 generations ago ranges between 2.5 cM and 0.94 centiMorgans (see Methods for details). Taking into account that multi-path IBD tracts will be on average longer[29], the IBD sharing signal at 1–5 cM detected between the western Turkic peoples and the SSM area populations may be due to historical Turkic and Mongolic expansions from the SSM area. It is possible to approximately outline the age of common ancestors directly from the distribution of shared IBD tracts [28], but such an inference would be too coarse for our purposes. Here we use two different methods implemented in ALDER [30] and SPCO [31] to infer the age of Siberian/Asian admixture among Turkic peoples. The admixture dates for all the analyzed Turkic peoples (Figure 5) fell within the historical time frame (5th–17th century) that overlaps with the period of nomadic migrations triggered by Turkic (6th–16th centuries CE) and Mongol expansions (13th century) [2,3]. However, individual admixture dates estimated using the two methods overlap only partially and were discordant for most populations (Figure 5). Therefore, we simulated a series of admixture events spanning a target historical period and compared how the two methods performed (see Material and Methods for details). The dates inferred by ALDER tended to be closer to simulated true values, while SPCO consistently estimated older dates (Figure 6). Importantly, the SPCO-inferred dates for our real dataset (Figure 5) also tended to be older, and we therefore suspect bias in our SPCO estimates. From here onward we discuss only ALDER-inferred dates.

**Figure 5.**
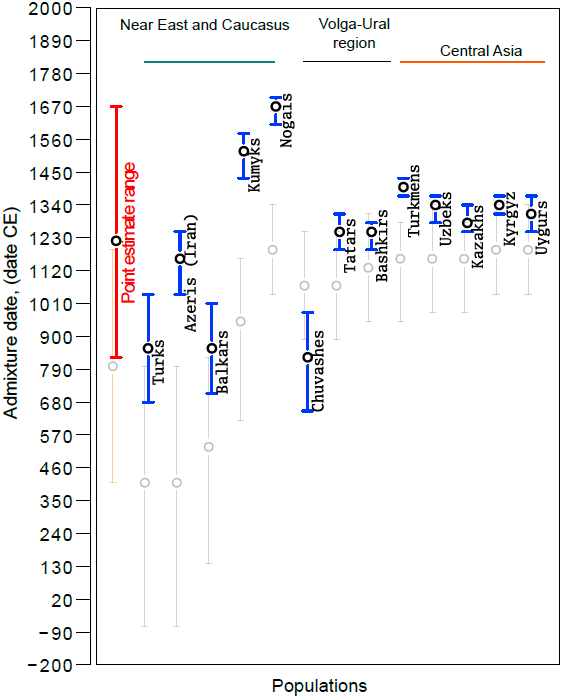
**Admixture dates for Turkic-speaking populations on an absolute date scale.** Blue circles show ALDER-inferred point estimates and error bars indicate 95% confidence intervals. Gray circles show SPCO-inferred point estimates and error bars in gray indicate 95% confidence intervals. The red bar shows the point estimate range (inferred using ALDER) across all the analyzed samples and the orange bar shows the same for SPCO-inferred dates. Admixture dates before Common Era (CE) are shown with a negative sign.

**Figure 6.**
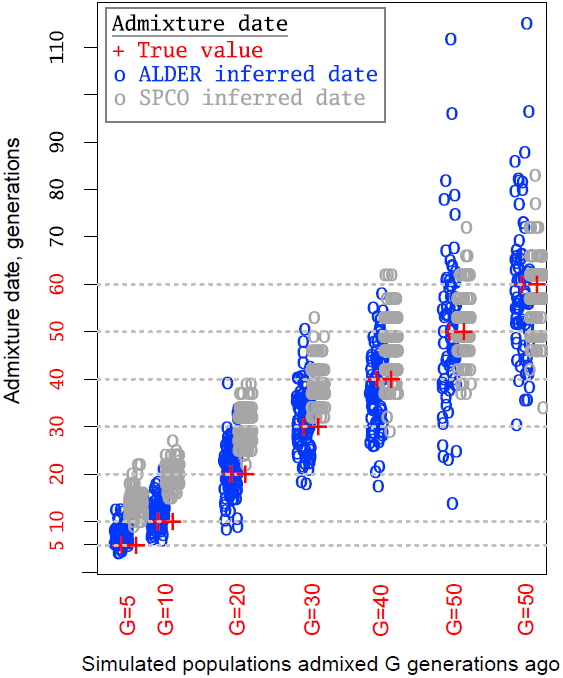
**Admixture dates for simulated populations.** Simulated populations were generated by mixing two ancestral populations G generations ago as described in the M&M section. We repeated each admixture scenario 120 times and analyzed with two admixture dating methods: ALDER and SPCO. Circles represent admixture dates for one simulated population and circle color indicates the method of admixture inference as shown in the legend. Red “plus symbols” show the true admixture date.

Although we report a single admixture date for each population, we note that it is likely that the contemporary Turkic peoples were established through several migration waves [2-4,32]. Indeed, Turkic peoples closer to the SSM area (those from the Volga-Ural region and Central Asia) showed younger dates compared to more distant populations like Anatolian Turks, Iranian Azeris, and the North Caucasus Balkars. Only Nogais, the former steppe belt nomadic people, and Kumyks inhabiting northern slopes of the Caucasus stand out from this spatial pattern.

### Discussion

Our ADMIXTURE analysis (Figure 2) revealed that Turkic-speaking populations scattered across Eurasia tend to share most of their genetic ancestry with their current geographic non-Turkic neighbors. This is particularly obvious for Turkic peoples in Anatolia, Iran, the Caucasus, and Eastern Europe, but difficult to say about northeastern Siberian Turkic speakers, Yakuts and Dolgans, for which non-Turkic reference populations are absent. We also found that a higher proportion of Asian genetic components distinguishes the Turkic speakers all over West Eurasia from their immediate non-Turkic neighbors. These results support the model that expansion of the Turkic language family outside its presumed East Eurasian core area occurred primarily through language replacement, perhaps by the *elite dominance* scenario.

When the Turkic peoples settled across West Eurasia are compared with their non-Turkic neighbors, they demonstrate higher IBD sharing with populations from SSM and Northeast Siberia than expected due to isolation by distance (Figure 3). There are, however, two non-Siberian populations that also demonstrate high IBD sharing with the tested Turkic peoples, the Kalmyks and Maris. These exceptions need careful consideration in light of historical data and previously published studies. For example, the Mongol-speaking Kalmyks migrated into North Caucasus from Dzhungaria (the northwestern province of China at the Mongolian border) only in the 17th century [32], while Maris stand out from other geographic neighbors due to unusually high recent admixture with Bashkirs: they demonstrate higher IBD sharing with Bashkirs for all IBD tract length classes (from 1–2 cM up to 11–12 cM) compared to other populations in the region (*p* < 0.05). This might be explained by the fact that we collected Maris samples in the Republic of Bashkortostan, where they seemingly intermarried with Bashkirs to some extent. Finally, some of the Siberian populations are in fact migrants in their current locations. For example, Yakuts, Evenkis, and Dolgans largely stem from the Lake Baikal region, which is essentially the SSM area [22]. It turns out that most of the populations showing a high signal of IBD sharing with the western Turkic populations originated from the SSM area or had admixture with one of the tested Turkic populations. The only exception is the Nganasans; they demonstrate unusually high IBD sharing with both western Turkic peoples (Figure 3) and randomly chosen non-Turkic populations (Figure S3). Taking into account that SSM area populations (Tuvans, Mongols, and Buryats) can be reliably considered indigenous to their locations, and that other Siberian and non-Siberian populations (demonstrating high IBD sharing with western Turkic peoples in Figure 3) all have SSM origins, we suggest that ancestral populations from this area contributed recent gene flow into western Turkic peoples. This recent gene flow, however, was not restricted to Turkic populations since our IBD sharing analysis revealed relatively high amounts of IBD sharing with the SSM populations for some non-Turkic peoples, such as Lezgins, North Ossetians, and Maris. This observation demonstrates that the inferred gene flow from the SSM area also contributed to the gene pool of non-Turkic peoples, but the stronger interaction (reflected in higher IBD sharing) with migrant SSM ancestors probably drove Turkicization, because modern Turkic peoples consistently show higher IBD sharing compared to their geographic neighbors. We performed a permutation test for each western Turkic population and the observed excess of IBD sharing with the SSM area populations was statistically significant (Figure 4 and Figure S4).

Another important outcome of our IBD sharing analysis is the finding that two of the three SSM populations that we consider “source populations” or modern proxies for source populations are both Mongolic-speaking. This observation can be explained in several ways. For example, one may surmise that the Mongol conquests, starting in the 13th century, were accompanied by their demographic expansion over the territories already occupied, in part, by Turkic speakers, and this led to admixture between Turkic and Mongolic speakers. Alternatively, it is also probable that the ancestors of Turkic and Mongolic tribes stem from the same or nearly the same area and underwent numerous episodes of admixture before their respective expansions. The latter explanation is indirectly testified by a complex, long-lasting stratigraphy of Mongolian loan words in Turkic languages and *vice versa* [33]. The first explanation is unlikely from a historical perspective since although Mongolic conquests were launched by Genghis Khan troops in the early 13th century, it is well known that they did not involve massive re-settlements of Mongols over the conquered territories. Instead, the Mongol war machine was progressively augmented by various Turkic tribes as they expanded, and in this way Turkic peoples eventually reinforced their expansion over the Eurasian steppe and beyond [34]. Therefore, we prefer the second explanation, although we cannot entirely exclude the Mongol contribution, especially in light of admixture dates that overlap with the Mongol expansion period.

Finally, our IBD sharing analysis suggested that the SSM area is the source of recent gene flow. This area is one of the hypothesized homelands for Turkic peoples and linguistically related Mongols. While the presence of the Mongol empire over this territory is well recorded, historical sources alone are insufficient to unambiguously associate this area with the Turkic homeland for several reasons: some of the Turkic groups speaking the Oghuric branch of Turkic were attested westerly in the Ponto-Caspian steppes in the mid-late 5th century CE. This is geographically distant from the SSM area, and temporarily much earlier than the Göktürk Empire was established in the SSM area. Thus, our study provides the first genetic evidence supporting one of the previously hypothesized IAHs to be near Mongolia and South Siberia.

The gene flow from the SSM area that we inferred based on our IBD sharing analysis should also be detected using an alternative approach such as ALDER, which is based on the analysis of linkage disequilibrium (LD) patterns due to admixture. Using the ALDER method, we tested all possible combinations of reference populations in our dataset. LD decay patterns observed among western Turkic populations were consistent with admixture between West Eurasian and East Asian/Siberian populations (see detected reference populations in Table S3). Admixture dating with the set of East Asian/Siberian populations (Table S3) inferred admixture events ranging between 816 CE for Chuvashes and 1657 CE for Nogais. We chose these reference populations based on the highest LD curve amplitudes, as suggested by the authors of the method. It is notable that all the SSM populations that were inferred to be the source of SSM gene flow were filtered out by an ALDER pre-test procedure because of the shared admixture signal with the tested Turkic populations. Indeed, as we show, SSM populations and the two Northeastern Siberian populations all demonstrated a statistically significant admixture signal between the same set of West Eurasian and East Asian populations as western Turkic peoples do (Table S3). Therefore, the set of reference populations reported in Table S3 that demonstrate the highest LD curve amplitude, in fact represent the set of closest possible reference populations that passed ALDER’s filtering procedure. This filter removes any reference population that shows shared admixture signal with the tested population. It was important for our study that the range of ALDER-inferred admixture dates overlaps with the major Turkic migrations and later Mongolic expansion (Figure 5), both of which are known to trigger nomadic migrations to Medieval Central Asia, the Middle East and Europe. In addition, when linguistic classification and regional context is taken into account, we found parallels with large-scale historical events. For example, the present-day Tatars, Bashkirs, Kazakhs, Uzbeks, and Kyrgyz span from the Volga basin to the Tien-Shan Mountains in Central Asia, yet (Figure 5) showed evidence of recent admixture ranging from the 13th to the 14th centuries. These peoples speak Turkic languages of the Kipchak-Karluk branch and their admixture ages postdate the presumed migrations of the ancestral Kipchak Turks from the Irtysh and Ob regions in the 11th century [32]. There are exceptions, like the Balkars, Kumyks, and Nogais in Northern Caucasus, who showed either earlier dates of admixture (8th century) or much later admixture between the 15th century (Kumyks) and 17th century (Nogais).

Chuvashes, the only extant Oghur speakers showed an older admixture date (9th century) than their Kipchak-speaking neighbors in the Volga region. According to historical sources, when the Onogur-Bolgar Empire (northern Black Sea steppes) fell apart in the 7th century, some of its remnants migrated northward along the right bank of the Volga river and established what later came to be known as Volga Bolgars, of which the first written knowledge appears in Muslim sources only around the end of the 9th century [35]. Thus, the admixture signal for Chuvashes is close to the supposed arrival time of Oghur speakers in the Volga region.

Differences in admixture dates for the three Oghuz speaking populations (Azeris, Turks, and Turkmens) were notable and their geographical locations suggest a possible explanation. Anatolian Turks and Azeris, whose Central Asian ancestors crossed the Iranian plateau and became largely inaccessible to subsequent gene flow with other Turkic speakers, both have evidence of earlier admixture events (12th and 9th centuries, respectively) than Turkmens. Turkmens, remaining in Central Asia, showed considerably more recent admixture dating to the 14th century, consistent with other Central Asian Turkic populations and most likely due to admixture with more recent, perhaps recurrent, waves of migrants in the region from SSM.

In summary, our collection of samples, which covered the full extent of the current distribution of Turkic peoples, shows that most Turkic peoples share considerable proportion of their genome with their geographic neighbors, supporting the *elite dominance* model for Turkic language dispersal. We also showed that almost all the western Turkic peoples retained in their genome shared ancestry that we trace back to the SSM region. In this way, we provide genetic evidence for the Inner Asian Homeland (IAH) of the pioneer carriers of Turkic language, hypothesized earlier by others on the basis of historical data. Furthermore, because Turkic peoples have preserved SSM ancestry tracts in their genomes, we were able to perform admixture dating and the estimated dates are in good agreement with the historical period of Turkic migrations and overlapping Mongols expansion. Finally, much remains to be learned about the demographic consequences of this complex historical event and further studies will allow the disentangling of multiple signals of admixture in the human genome and fine scale mapping of the geographic origins of individual chromosomal tracts.

## Material and Methods

### Ethics Statement

All subjects signed personal informed consents and ethical committees of the institutions involved approved the study.

### Samples, Genotyping, and Quality Control

In total, 322 individuals from 38 populations were genotyped on different Illumina SNP arrays (all targeting > 500,000 SNPs) according to manufacturers’ specifications. Our data was combined with published data from Li et al. [19], Rasmussen et al. [20], Behar et al. [21], Yunusbayev et al. [14], Metspalu et al. [27], Fedorova et al. [22], Raghavan et al. [36], Behar et al. [37], and covered all the Turkic-speaking populations (373 individuals from 22 samples) from key regions across Eurasia and their geographic neighbors (see details about sample source in Table S1). Individuals with more than 1.5% missing genotypes were removed from the combined dataset. Only markers with a 97% genotyping rate and minor allele frequency (MAF) > 1% were retained. The absence of cryptic relatedness corresponding to first and second degree relatives in our dataset was confirmed using King [38]. The filtering steps resulted in a dataset of 1,444 individuals remaining for downstream analyses. It is important to note that in our dataset there are 312,524 SNPs that are common for Human1M-Duo and 650k, 610k, and 550k Illumina BeadChips. Different analyses have different requirements regarding marker density and we therefore prepared two datasets. For Admixture and ALDER analyses that require minimum background LD, LD pruning on the combined 1M-Duo and 650k, 610k, and 550k dataset was performed. The marker set was thinned by excluding SNPs in strong LD (pairwise genotypic correlation r^2^ > 0.4) in a window of 1,000 SNPs, sliding the window by 150 SNPs at a time. This resulted in a dataset of 174,187 SNPs. Another dataset with a dense marker set for IBD sharing and wavelet transform admixture dating analyses was prepared. For this, the 1M-Duo genotyped samples (Table S1) were excluded to increase the SNP overlap among remaining samples up to 515,841 markers. Genetic distances between SNPs in centiMorgans were incorporated from the genetic map generated by the HapMap project [39].

### Admixture analysis

We inferred population structure in our dataset using a model-based clustering method implemented in ADMIXTURE software. Because the ADMIXTURE algorithm expects SNPs to be unlinked, we used an LD pruned marker set of 174,187 SNPs. We ran ADMIXTURE assuming 3 to 14 (*K* = 3 to *K* = 14) genetic clusters or “ancestral populations” (see Figure S1) in 100 replicates and assessed convergence between individual runs. For low values of *K*, all runs arrive at the same or very similar log-likelihood scores (LLs), whereas runs using higher *K* values have more variable LLs. Relying on the low level of variation in LLs (LLs < 1) within a fraction (10%) of runs with the highest LLs, we assume that the global log-likelihood maximum was reached at *K* = 3 to *K* = 11 and *K* = 13 to *K* = 14 (Figure S6). ADMIXTURE provides an assessment of the “best” *K* by computing a cross-validation index (CV), which estimates the predictive accuracy of the model at a given *K*. In our setting clustering solutions at *K* = 5, 6, 7, and 8 showed better predictive power than other *K* values (Figure S7). In choosing which model(s) (*K*) to discuss further, we draw from the CV results and restrict ourselves to the *K*s that likely converged at the global LL maximum for the particular model. In addition, we acknowledge that clustering solutions at different *K*s may reflect the hierarchical nature of human population structure. In this study, we are interested in distinguishing regional groupings, like Northeast Asia and Europe, and possible admixture between such groups. In sum, we found that the clustering solution that met our selection criteria best was *K* = 8.

### IBD sharing analysis

We used the fastIBD algorithm [40] implemented in BEAGLE 3.3 software to detect extended chromosomal tracts (> 1 cM in length) that are IBD between pairs of individuals. We ran the fastIBD algorithm ten times with different random seeds and called IBD tracts using a modified post-processing tool ‘plus-process-fibd.py’. The original post-processing tool developed by the BEAGLE authors was modified by [28]. They added an algorithm that minimizes the number of spurious breaks and gaps introduced into long segments due to low marker density [28].

Patterns of IBD sharing between modern populations can bear information about historical events [28]. According to known history, the Turkic migrations took place roughly 600–1600 years ago. Assuming a human generation time of 30 years, the immigrant chromosomes left during Turkic migrations have passed through 600 / 30 = 20 and 1600 / 30 = 53 generations/meiosis. The mean length of a single-path IBD tract passed through ∼20–53 generations, is expected to be 100 cM / (2 * 20) = 2.5 cM and 100 cM / (2 * 53) = 0.94 cM [29]. We provide these estimates only as reference, and note that the true IBD tract length distribution for a given historical event is influenced by past demography. The fastIBD method that we use to search for IBD tracts has sufficient power (∼0.7– 0.9) to detect chromosomal tracts of 2–3 cM in length, and importantly, close to zero false discovery rate [40]. The power to detect IBD tracts of 1 cM in length varies from 0.2 to 0.5 depending on the chosen fastIBD score threshold. We used a fastIBD score threshold of 1e-10, which, in the trade-off between power-loss and minimizing false discovery rate favors the latter, keeping it close to zero. These parameter settings fit our purposes since we are interested in estimating the relative amount of IBD sharing between populations rather than the total amount of IBD sharing.

### Isolation-by-distance test

Chromosomal tracts that were IBD between two populations were first sorted into bins (classes) based on their length: 1–2, 2–3, 3–4, 4–5 cM, and then the total length was divided within each bin (class) by sample size to obtain the average IBD sharing for each population pair tested. From here onward, we refer to this statistic as IBD sharing. It was shown previously that IBD sharing between populations decays exponentially with distance between samples [28]. To test whether IBD sharing between populations in our dataset is consistent with isolation by distance, we first converted the IBD sharing statistic between populations into an IBD sharing distance using –ln(IBD sharing statistics). For each pair of populations, we then calculated geodesic distances in kilometers using the ‘‘distonearth’’ R function (Banerjee 2005). Geographic coordinates for populations in our dataset were calculated using the central point from multiple sampling locations, or when such coordinates were not available, using coordinates for the country center (where sample was collected). Geographic coordinates for HGDP populations were computed as a central point in a range of longitude and latitude values given in [41]. After obtaining matrixes of geodesic and IBD sharing distances between populations, values were standardized in each of these matrixes using the maximum values observed. The Mantel test was run on the obtained standardized distance matrixes using the mantel.rtest function in the ‘ade4’ R package[42].

### Identifying deviations from the isolation-by-distance pattern

Even if there is a statistically significant correlation between IBD sharing and geographic distance in the data, spatial patterns of recent ancestry between real populations are unlikely to meet isolation-by-distance expectations ideally. One way to detect the departure from the expected isolation-by-distance pattern is to compute parameters that describe the relationship between IBD sharing and distance in a population set where you do not expect any deviation. These parameters can then be used to compute the expected range of IBD sharing for a given pair of populations at a given distance and report deviations, if any. In our dataset, it was difficult to define a population subset that was completely devoid of samples with departures from the expected isolation-by-distance pattern. Therefore, a comparative approach was used, in which the IBD sharing pattern in our dataset with Turkic populations and without them was compared. Because departures from the isolation-by-distance pattern may already exist in the dataset without Turkic peoples, this test determines whether Turkic peoples have more extreme departure due to, for example, a long range migration from Asia, as suggested by our ADMIXTURE analysis. Therefore, the test dataset only included western Turkic populations from the Middle East, Eastern Europe, the Caucasus, and Central Asia. The overall goal is to find systematic (correlated) differences between these western Turkic populations and their geographic neighbors. The null hypothesis was that differences between western Turkic populations and their geographic neighbors are random. To perform this analysis, sets of geographic neighbors were defined for each of the 12 western Turkic populations (See Table S2; the same sets are used for permutation test described below; See Figure 4) and IBD sharing that these populations demonstrate with other samples in our dataset was computed. A total of 63 Eurasian populations were included in our dataset (excluding 12 western Turkic populations). Thus, for each set of geographic neighbors, a vector of 63 ordered IBD sharing values with other populations in the dataset was obtained, including self-comparisons. For each of the western Turkic populations, the same vector of IBD sharing values was computed (the Turkic population versus 63 Eurasian populations). After performing element-wise subtraction of values in the “vector for geographic neighbors” from values in the “vector for Turkic population”, differences in IBD sharing were obtained. Provided that western Turkic populations have no systematic difference with their geographic neighbors, differences are expected to occur at random; that is, a vector of 63 random values is expected (positive when greater than average, negative when lower than average, and zero when equal to average). By contrast, if some of the 63 populations in the dataset, for example, Romanians, systematically demonstrate higher IBD sharing with all the tested Turkic populations, positive values are expected for Romanians. Such nonrandom signals can be detected by computing an element-wise sum over all the vectors (for all the 12 Turkic populations) containing differences. If they are random, there will be no accumulation. We call this procedure “subtraction and accumulation” throughout the text. Element-wise addition of vectors would accumulate positive IBD sharing values for Romanians (versus 12 Turkic populations) and when such values are plotted on a geographic map, a very high signal of (due to accumulation) IBD sharing should be observed for Romanians, compared to the other 63 samples. See the schematic representation of this “subtraction and accumulation” procedure in Figure S2. An accumulated value (IBD sharing signal) for a given population was considered high when it exceeded the 0.90 sample quantile point. Finally, this “subtraction and accumulation” procedure was repeated multiple times by replacing each of the 12 Turkic populations with randomly chosen non-Turkic neighbors from respective sets of geographic neighbors (see Figure S3 for results). Doing so demonstrates the kind of results expected when the “subtraction and accumulation” procedure is done with population sets that do not have systematic differences in IBD sharing.

### Permutation test

A permutation test was designed to verify whether the excess of IBD sharing that western Turkic populations demonstrate with SSM populations is statistically significant. IBD sharing (as described previously) between a western Turkic-speaking population and each of the three SSM populations (Tuvans, Buryats, Mongols) that show high accumulated IBD sharing was calculated. A permutation procedure was then used to test whether observed excess of IBD sharing that a given Turkic population demonstrates can be expected by chance among its non-Turkic neighbors. For each Turkic population, their geographic neighbors were pooled and 10,000 random samples of the same size were generated (as for Turkic population tested). For each random sample, IBD sharing with the three SSM populations and Evenkis was calculated. Obtained IBD sharing statistics from permuted samples were compared to that of Turkic populations from the same region and the number of tests showing equal or higher values was divided by the total number of permutations to obtain a *p*-value.

### Admixture dating using ALDER

We used ALDER[30] to test whether the gene flow from SSM area suggested by our IBD sharing analysis left detectable trace in LD pattern among the Turkic populations, and date this admixture signal. ALDER has a functionality to perform a statistical test for the presence of admixture and dating admixture signal. We tested all possible combinations of populations in our dataset to be reference group for Turkic populations and report a pair that successfully passed all the pre-test steps and has significant p-value for admixture. By choosing the pair of reference populations with the highest LD curve amplitude we report ALDER-inferred admixture date for the admixed population.

### Dating admixture using wavelet transform method

For each Turkic-speaking population, two parental populations were selected, so that one represented local ancestry and another represented immigrant SSM ancestry. Based on our IBD sharing analysis, Tuvans were used to represent the SSM parental population for all Turkic-speaking populations. Alternative SSM ancestors were also tested (See Table S4). The parental population that represented local ancestry was chosen among geographic neighbors. The wavelet transform method as implemented in the StepPCO software was used [31]. This software projects an admixed population, in this case the Turkic-speaking population, along a principal component separating the two chosen ancestors, the Tuvans, representing the SSM ancestor and a geographic neighbor (see Table S4). At this step, PC coordinates for each projected Turkic population were used to infer the proportion of admixture contributed by each parental population. Given that x1 and x2 are the average PC coordinates for the first and second parental populations and x3 is the average PC coordinate for the admixed population, the proportion of ancestry contributed by the first parental population is α = (x2 - x3) / (x2 - x1) [43]. Accordingly, the proportion of ancestry contributed by the second parental population is (1 - α). Admixture proportions inferred at this stage were used later to find a matching simulated dataset with the same admixture proportion. The StepPCO software infers which chromosomal tracts have local or immigrant SSM ancestry and uses observed length distributions of the ancestry tracts to compute a genome-wide wavelet transform (WT) coefficient [31]. These WT coefficients for each Turkic-speaking population are compared with those of simulated samples that have matching admixture proportions and for which the admixture history is known. Samples with known admixture history were generated using a forward simulation in the SPCO software. In this forward simulation, a population with effective population size of 1,000 individuals at time T_0_ receives migrants from another population and grows for 300 generations until it reaches an effective population size of 10,000 individuals. We modeled different amounts of migrants, replacing 5%, 10%, 15%, 20%, …, 75% of the recipient population. Each admixture scenario was repeated 100 times and a random sample of 20 individuals was drawn to compute WT coefficients. The WT coefficients from these 100 independent runs were used to construct a 95% confidence interval for a given admixture scenario. Simulated WT coefficients with 95% confidence interval were plotted against the known number of generations since admixture (Figure S5). Admixture time for tested populations was obtained by comparing point estimates with the curve of simulated WT coefficients.

### Coalescent simulations

To test how different dating methods recover signals of admixture known to have occurred during historical periods, gene flow events occurring 5, 10, 20, 30, 40, 50, and 60 generations ago were simulated in 120 repetitions. The MaCS coalescent simulator [44] was used to generate a series of historical gene flow events between two parental populations whose demographic parameters imitate that of Asian and European populations. Demographic parameters (split time, growth rate, and bottlenecks) for the two simulated parental populations were taken from the study by Schaffner et al. [45]. In this study, a series of population genetic statistics were used to fit the demographic histories of simulated populations to those observed for African, Asian, and European populations. Here, these best-fitting demographic parameters were used to simulate samples that imitate sequences drawn from Asian and European populations. Each “historical time” gene flow event represents a mixture of the two parental populations (Asian and European) at equal proportions. To mimic the variation in recombination rate observed in real populations, sequences (250 Mb in length) were simulated using the recombination (cM/Mb) mappings of chromosome 1 from HapMap project phase 2 [39]. From each simulated admixed population a sample of 30 sequences were drawn to construct 15 genotypes that were then subjected to the same data preparation and quality control steps as for real data during admixture dating. Two admixture dating approaches were then applied to our simulated datasets: ancestry block distribution based (Wavelet transform method) and weighted LD curve based (ALDER).

## Acknowledgments

We thank the individuals who provided DNA samples for this study. We thank the Estonian Biocentre core facility and Estonian Genome Center of the University of Tartu (EGC-UT) technical personnel, especially Tuuli Reisberg, Viljo Soo, and S. Smit for sample handling and conducting the autosomal genotyping. We are thankful to Georgi Hudjashov for preparing autosomal data, and to Brian Browning and Sharon Browning for their helpful discussions on our preliminary IBD sharing results.

## Accession numbers

All Illumina genotyping data can be accessed through our website at http://evolbio.ut.ee/.

## Supporting Information

**Figure S1.** Population structure inferred using a structure-like approach assembled in ADMIXTURE (at *K* = 2 to *K*11, *K*13, and *K*14). Each individual is represented by a vertical (100%) stacked column indicating the proportions of ancestry in *K* constructed ancestral populations. The models (*K*) shown here likely each converged to a global likelihood maximum as > 10% of runs (100 replicates in total for each *K*) with the highest log-likelihood (LL) converged to essentially to the same solution with a log-likelihood difference of > 1 LL units. We plotted the runs with the highest LL at each *K*. (PDF)

**Figure S2.** Schematic representation of the “subtraction-accumulation” analysis aimed at detecting correlated signals of IBD sharing that different western Turkic populations show with others in the dataset. Panels A, B, C, and D show sequence of steps in the analysis (PDF)

**Figure S3.** Populations with high and correlated signals of IBD sharing with randomly selected geographic neighbors for western Turkic peoples. Circle position corresponds to population location. Circle color indicates the amount of excess IBD sharing (shown in the Legend) that this population shares with all 12 randomly selected geographic neighbors. Populations with IBD sharing exceeding the 0.90 quantile are shown with the “plus symbol”. Panels from A–J show IBD sharing signals for different randomly selected combinations of geographic neighbors. All the results are based on IBD tracts of 1–2 cM. (PDF)

**Figure S4.** Pairwise IBD sharing summary binned by segment size. Panels A, B, C, and D show results for different segment size bins (2-3, 3-4, 4-5, 5-6 cM). For each population ordered along the x–axis, IBD sharing is computed with four populations (Tuvans, Buryats, Mongols (Mongolia), and Evenkis) from the SSM area. Each Turkic-speaking population (shown in red) is grouped with its respective geographic neighbors using parentheses. The grouped geographic neighbors were pooled and used to perform a permutation test as described in the M&M section. The red number under the Turkic population name shows how many SSM populations demonstrate a statistically significant excess of IBD sharing with a given Turkic population. Overlapping parentheses show Turkic-speaking populations with shared geographic neighbors. (PDF)

**Figure S5.** Admixture time estimates using the SPCO method. The blue curve shows the relationship between the WT coefficient and time since admixture in a growing population. Each curve summarizes the outcome from 100 forward simulations. Thus, the bold curve shows the average WT coefficient over 100 simulations and the blue shaded area shows the 95% confidence interval. Horizontal lines in red, green, and blue show point estimates of the WT coefficient for different Turkic-speaking populations. The intersection point between the horizontal line and the blue curve gives the admixture time estimate, shown with dashed vertical lines. (PDF)

**Figure S6.** The maximum difference in log likelihood (LL) scores in fractions (0.05, 0.1, and 0.2) of ADMIXTURE runs with the highest LL scores. For clarity, the y-axis is shown in two sections. (PDF)

**Figure S7.** A boxplot of cross-validation errors of the ADMIXTURE runs in 100 replicates at K values 2 through 14. (PDF)

**Table S1.** Population samples and genotype sources. (XLS)

**Table S2.** Population samples used in the “subtraction-accumulation” IBD sharing analysis. (XLS)

**Table S3.** Admixture dates based on weighted LD statistics. Admixture dates in years were estimated as the number of generations multiplied by 30 years. Absolute dates or Calendar dates were estimated as 2000 - (30 * generations). (XLS)

**Table S4.** Admixture dates based on the wavelet transform (SPCO) method. (XLS)

